# Plant spectral diversity from high-resolution multispectral imagery detects functional diversity patterns in coastal dune communities

**DOI:** 10.1101/2023.02.07.527269

**Authors:** Eleonora Beccari, Carlos Pérez Carmona, Enrico Tordoni, Francesco Petruzzellis, Davide Martinucci, Giulia Casagrande, Nicola Pavanetto, Duccio Rocchini, Marco D’Antraccoli, Daniela Ciccarelli, Giovanni Bacaro

## Abstract

- Remote sensing is a fundamental tool to monitor biodiversity over large spatial extents. However, it is still not clear whether spectral diversity (SD - variation of spectral response across a set of pixels) may represent a fast and reliable proxy for different biodiversity facets such as taxonomic (TD) and functional diversity (FD) across different spatial scales.
- We used fine resolution (3 cm) multispectral imagery on coastal dune communities in Italy to explore SD patterns across spatial scales and assess SD relationships with TD and FD along the environmental gradient.
- We measured TD as species richness, while SD and FD were computed using probability densities functions based on pixels and species position in multivariate spaces based on pixel values and traits, respectively. We assessed how SD is related to TD and FD, we compared SD and FD patterns in multivariate space occupation, and we explored diversity patterns across spatial scales using additive partitioning (i.e., plot, transect, and study area).
- We found a strong correspondence between the patterns of occupation of the functional and spectral spaces and significant relationships were found along the environmental gradient. TD showed no significant relationships with SD. However, TD and SD showed higher variation at broader scale while most of FD variation occurred at plot level.
- By measuring FD and SD with a common methodological framework, we demonstrate the potential of SD in approximating functional patterns in plant communities. We show that SD can retrieve information about FD at very small scale, which would otherwise require very intensive sampling efforts. Overall, we show that SD retrieved using high resolution images is able to capture different aspects of FD, so that the occupation of the spectral space is analogous to the occupation of the functional space. Studying the occupation of both spectral and functional space brings a more comprehensive understanding of the factors that influence the distribution and abundance of plant species across environmental gradients.

## 1. Introduction

Global change is dramatically affecting global biodiversity (Butchart *et al*., 2010; Winkler *et al*., 2021), with no signs to decrease in the near future (Díaz et *al*., 2019; Trisos et *al*., 2020). To keep up with these changes, scientists have proposed to consider fast and repeatable measures of all facets of biodiversity over large extents (McGill *et al*., 2015; Jetz *et al*., 2016). In this context, remote sensing emerges as the most comprehensive and convenient tool for handling multiple biodiversity-related questions (Jetz *et al*., 2016) and is particularly promising in detecting different facets of plant diversity (Yannelli *et al*., 2022). Yet, major gaps remain in the application of remote sensing to detect biodiversity patterns across different spatial scales (Wang & Gamon, 2019).

Remotely sensed images can be used to detect the morphological, physiological, and chemical structures of vegetation (Ustin *et al*., 2009; Ollinger, 2011). Because differences in vegetation phenotypic characteristics correspond to variations in spectral band values (Spectral Variation Hypothesis; Palmer et *al*., 2002; Rocchini *et al*., 2004), spatio-temporal variations in spectral bands (i.e. spectral diversity, SD) can be considered an indicator of spatio-temporal variability of vegetation. This makes SD a cost- and time-efficient proxy for different diversity facets (Surrogacy hypothesis; Gamon *et al*., 2020; Wang & Gamon, 2019), such as taxonomic diversity (TD, e.g. Conti *et al*., 2021; Marzialetti *et al*., 2021) and functional diversity (FD, e.g. Frye et *al*., 2021; Zhao et *al*., 2021).

Several studies have explored the effectiveness of SD in approximating TD (Torresani *et al*., 2019; Conti *et al*., 2021), showing a strong context-dependency of the relationships between remotely sensed plant spectrum and species identity (Schmidtlein & Fassnacht, 2017; Fassnacht et *al*., 2022). Specifically, both image resolution (i.e., pixel and spectral resolution, Gamon *et al*., 2020) and vegetation characteristics (e.g., vegetation type and size) influence spectral variability, producing contrasting patterns between SD and TD that are sensor- and ecosystem-dependent (Schmidtlein & Fassnacht, 2017; Conti *et al*., 2021; Fassnacht *et al*., 2022).

Spectral discrimination of plant species is based on plant attributes, which are products of the set of plant traits that can be remotely detected (Fassnacht *et al*., 2022). Because of this, plants’ spectral properties have a strong link with plant traits (Ustin & Gamon, 2010; Homolová *et al*., 2013), as a consequence a stronger relationship is expected with FD rather than with TD (Conti *et al*., 2021). In this context, hyperspectral imaging is efficient in discriminating trait variation among communities (Frye *et al*., 2021), and has shown a great potential in depicting ecological processes (Schweiger *et al*., 2021). However, a widespread use of hyperspectral sensors in research is still limited due to their high costs coupled with computational-intensive management (Adão *et al*., 2017; Rossi *et al*., 2022). Recent studies have shown that images with lower spectral resolution (i.e. multispectral), which are a feasible alternative, are also capable to detect not only plant traits across different ecosystems and pixel resolutions (Aguirre-Gutiérrez *et al*., 2021; Thomson *et al*., 2021) but have also the potential to detect changes in the functional structure of the communities (Hauser *et al*., 2021; Helfenstein *et al*., 2022).

FD and SD are complex by nature: they derive from the combination of multiple traits and bands, respectively, and reflect differences among species coexisting in an assemblage. Therefore, both FD and SD should be addressed including all aspects of their variation. As a result, to efficiently depict functional patterns and processes in a given assemblage, it is pivotal to address FD in an holistic way (Mason *et al*., 2005). However, researchers generally use synthetic measurements of single aspects of functional variation, such as the mean (community weighted mean, Ricotta & Moretti, 2011), the variance (Rao’s Q, de Bello *et al*., 2016) or the combination of size and location of species within a trait space (convex hull, Villéger *et al*., 2008) that can overlook the strong constraints and coordination patterns underlying the functional structure of plant assemblages (Díaz *et al*., 2016; Carmona *et al*., 2021a). Conversely, addressing all aspects of FD simultaneously in a multivariate trait space can provide a direct access to community and ecosystem changes (Díaz *et al*., 2016; Joswig *et al*., 2022). The trait probability density approach (TPD; Carmona *et al*., 2016) is appropriate for this task, since it reflects the probabilistic distribution of species in a trait space (Carmona *et al*., 2021a; Rodríguez-Alarcón *et al*., 2022). Similarly to FD, approaches measuring SD generally rely on synthetic measurements of single aspects of its variation, such as ratios of single bands (e.g. NDVI, Torresani *et al*., 2019), variance (Rao’s Q, (Rocchini *et al*., 2017) or the combination of size and location of pixels within a spectral space (convex hull, Gholizadeh *et al*., 2018). Expanding the TPD approach to represent SD would not only allow us to consider it in a holistic way including simultaneously all aspects of spectral variation across multiple spectral bands, but also provide a common toolbox to better compare SD and FD across different spatial scales.

In this study, we combined fine-scale (3 cm) multispectral airborne remotely sensed images coupled with an intense field sampling survey, to evaluate the ability of remote sensing in retrieving biodiversity patterns in Mediterranean coastal dunes ecosystems. Coastal dune ecosystems are defined by a strong sea-inland gradient, characterized by high taxonomical turnover and increasing species richness proceeding landward (Tordoni *et al*., 2018, 2021). From a functional perspective, coastal dune communities are defined by different vegetative growth forms along the sea-inland gradient (i.e., herbaceous and woody) with generally higher FD closer to the sea where environmental conditions are harsher and species richness is low (Carboni *et al*., 2013).

We addressed SD by calculating a multidimensional space based on the spectra of pixels and estimated SD using the TPD framework originally developed by Carmona et al. (2016) in a FD context. By using the TPD framework, we can derive functional and spectral structures (i.e., patterns of organization of the species — or pixels — within the functional or spectral space) that can be used to compare and quantify patterns in space occupation. In doing so, we produce comparable estimations of all aspects of FD and SD allowing us to assess the SD-FD relationship in a way that goes beyond the traditional comparisons of univariate indicators of variation. We used this framework (i) to test whether SD can approximate patterns in plant diversity (both TD and FD), (ii) to detect how TD, FD, and SD patterns are coordinated along the sea-inland gradient, and (iii) to explore how TD, FD, and SD partition across different spatial scales (plot, transect, study area). Given the fine pixel resolution, we hypothesize that SD will better capture FD than TD because it is directly related to species’ phenotypes rather than their taxonomic identities. Consequently, we hypothesize a strong correspondence between functional and spectral structures, which should be further corroborated by a high positive correlation between functional and spectral dissimilarities. Further, if patterns in space occupation are strongly coordinated, we also expect that the amount of space occupied by each plot and transect (i.e. richness, Mason *et al*., 2005) in the functional and spectral space should be positively correlated and this correlation should be reflected along the sea-inland. Finally, we expect that the correspondence between FD and SD across plot and transects will produce similar diversity partitioning across spatial scales.

## 2. Material and Methods

### 2.1 Study area and sampling design

Coastal dune ecosystems are characterized by steep ecological gradients (Tordoni *et al*., 2021), producing marked vegetation zonation arrange from sea to inland (Acosta *et al*., 2007; Tordoni *et al*., 2018). Closer to the sea (i.e., foredunes), plant species experience harsher abiotic conditions that limit the establishment to extremely specialized herbaceous species (Acosta *et al*., 2008). In the landward part of the beach (i.e., fixed dunes), the harshness of these conditions decreases allowing plant communities to become richer in species and growth forms (e.g., small shrubs of *Juniperus* spp., Acosta *et al*., 2008).

Fieldwork was performed in May-June 2019 within the protected coastal sand dune habitats of “Migliarino-San Rossore-Massaciuccoli” Regional Park (Italy). The study area belongs to the Natura 2000 network (Directive 92/43/EEC) and includes the Special Area of Conservation “Dune Litoranee di Torre del Lago” (IT5170001, centroid coordinates 10.253889E, 43.828611N). The sampling design for the collection of plant data was based on the selection of a squared grid of 500 ×500 m overlaying the whole study area. Within each cell of the grid, one transect was randomly selected and placed from sea-landwards for a total of 6 transects sampled in the whole study area. According to the dune morphology and extension, transects’ length ranged between 172 and 208 m encompassing a set of contiguous squared plots of 16 m^2^ each, in which we assessed the occurrences and abundance of vascular plant species (measured as percentage visual cover). We sampled a total of 288 plots in the six transects. Plant nomenclature was standardized according to Bartolucci *et al*. (2018) and Galasso *et al*. (2018).

### 2.2 Functional traits measurement and remotely sensed image processing

Leaf samples for functional traits measurements were randomly collected within transects trying to maximize interspecific trait variation. To this aim, we sampled 2 to 4 individuals for each species both in foredunes and fixed dunes. In total, functional traits were measured on 42 out of 75 recorded species, accounting for 97.4% of the total plant coverage. Following Petruzzellis et al. (2021), we measured cost-related, hydraulic and leaf vein traits associated to the “Leaf Economics Spectrum” (LES; Wright *et al*., 2004) and to the “flux trait network” proposed by Sack et al. (2013). Specifically, we selected the following functional traits: Specific Leaf Area (SLA, mm^2^/mg); Leaf Dry Matter Content (LDMC, mg/g); Minor Vein Length per unit Area (VLA_minor_, mm/mm^2^); Water potential at turgor loss point (Ψ_tlp_, - MPa). Detailed procedure for functional traits measurements is reported in supplementary material.

Remotely sensed image acquisition was performed immediately before vegetation sampling. We conducted Unmanned Aerial Vehicle survey using a MicaSense RedEdge 3© multispectral camera (MicaSense, Seattle, WA). We acquired one image per transect at 3 cm pixel resolution using five multispectral bands: blue (center wavelength: 475 nm), green (560 nm), red (668 nm), Near Infra-Red (NIR, 840 nm), and Red Edge (RE, 717 nm). Considering the patchy vegetation and the bare sand defining coastal dune ecosystem we decided to remove all pixels not related to vegetation. We performed an unsupervised linear spectral unmixing process (Settle & Drake, 1993) which has been previously used to separating pure and mixed pixels in coastal dunes (Lucas *et al*., 2002). For each transect, we used only pixels containing a minimum of 60% of vegetation for successive analysis. Spectral cleaning procedure produced a total of ca. 1,5 million pixels used for subsequent analysis. It is worth mentioning that the whole spectral cleaning procedure may have led to lose small or hidden (i.e., covered by sand) species, particularly in plots closer to the sea where smaller and more scattered individuals were present. See supplementary materials for a full description of images processing.

### 2.3 Estimation of the functional and spectral spaces

Mapping species position in multidimensional spaces based on trait information (i.e., functional space) allows summarizing species’ ecological strategies through a reduction of their dimensionality (Díaz *et al*., 2016). We defined a functional space using a Principal Component Analysis (PCA) considering mean functional traits values for each species (*n* = 42). Using Horn’s Parallel Analysis implemented in the *‘paran’* R package (Dinno, 2018), we retained two axes, which accounted for 76.8% of trait variation.

Similarly, mapping pixels position in a multidimensional space based on band values, allows to summarize the reflectance spectrum of pixels conveying information on all bands simultaneously (Conti *et al*., 2021; Rocchini *et al*., 2021). We first removed outliers from each of the 5 spectral bands by deleting pixels laying outside the 95% of the data distribution to reduce spectral aberration. Then, we performed a PCA using multispectral band values for every single pixel of the study area (*n* = 1,470,416), obtaining a multivariate space (i.e., spectral space). We retained the first two components, accounting for 89.3% of pixel variation. In this approach, we assumed that single pixels are equivalent to “spectral individuals”, while spectral bands can be considered as the equivalent of “spectral traits”. Under these assumptions, the scores of each pixel in the retained PCA axes reflect the position of the pixel in the spectral space, in the same way that the scores of species in the PCA based on traits reflect the position of the species in the functional space. In order to produce reliable Trait Probability Density estimations (see following paragraph), only plots with at least 5 pixels were kept for subsequent analyses, for a total of 270 plots (out of 288 plots sampled) with 5,446 ± 2,836 pixels per plot (mean ± std. dev.).

### 2.4 Estimation of functional and spectral structures and diversity metrics

We estimated TD at the different hierarchical scales of the study: i.e., within plots (α– diversity), among the plots from the same transect (β–diversity), and at the whole study area (γ–diversity). α–TD was measured as species richness, whereas β–TD was calculated as pairwise Sorensen dissimilarity, using the function *beta.pair* available in R package *‘betapart’* (Baselga *et al*., 2021).

We then derived functional and spectral structures for each plot (i.e., the patterns of organization of the species - or pixels - in a plot within the functional or spectral space) using the trait probability density framework (TPD; Carmona *et al*., 2016). TPD functions are probability density functions so that they reflect the probabilistic distribution of points (in our case species or pixels) in a given space (functional or spectral, respectively). We estimated functional and spectral TPD functions for each plot using the *‘TPD’* R package (Carmona *et al*., 2019). Then, we estimated different aspects of functional and spectral diversity within plots (α– diversity) and among the plots from the same transect (β–diversity). In particular, within-plots we estimated functional (and spectral) richness which reflects the amount of the functional (and spectral) space that is occupied by any given plot (Carmona *et al*., 2016). β–diversity was expressed as the dissimilarity between pairs of plots from the same transect. Plot level functional (β–FD) and spectral dissimilarities (β-SD) were calculated as 1 – overlap of the corresponding TPD functions (Carmona *et al*., 2019). TPD-based dissimilarities consider both differences in boundaries and differences in how densely the functional (or spectral) space is occupied by each of the compared plots, thus providing an indication of dissimilarity between communities that encompasses simultaneously all aspects of functional (or spectral) variation. We further decomposed dissimilarity into two complementary components, namely turnover and nestedness: turnover quantifies up to what point the functional (and spectral) differences between plots are because the plots occupy exclusive regions of the space, whereas nestedness reflects how differently the two plots occupy the parts of the functional (and spectral) space that they share. Finally, TPD functions of all plots contained in each transect were additively aggregated to produce transect-level TPD functions, and the same methods and metrics described above were estimated at the transect level.

### 2.5 Correspondence among functional structure, spectral structure, and species composition

To test the hypothesis that multispectral SD can approximate FD better than TD, we explored up to what point spectral structures are able to reflect functional structures and species richness between plots. First, we assessed the correlation between functional, spectral, and taxonomical dissimilarities among pairs of plots. To do this, we performed Mantel tests (Legendre & Legendre, 2012), using Spearman correlation coefficient (ρ) and 999 randomizations. Then, to test if correlations between plot dissimilarities were actually driven by their functional and taxonomical covariation, respectively, we performed Partial Mantel tests (Legendre & Legendre, 2012) between β–SD and its functional and taxonomic counterparts while controlling for the third component. Both tests were performed using the functions *mantel* and *partial.mantel* in the *‘vegan’* R package (Oksanen *et al*., 2020). The same set of analysis was also repeated within each transect. Finally, we explored the relationships between β–SD and both β–FD and β–TD using major axis regressions for each transect. Major axis regression models do not assume causality between variables and account for errors in variable estimation (Legendre & Legendre, 2012), allowing us to accurately describe relationships between SD and biodiversity. All models were performed using *lmodel2* function of the *‘lmodel2’* R package (Legendre, 2018).

Addressing plot pairwise dissimilarities allow us to include all aspects of functional and spectral variation by directly addressing the probabilistic distributions of species and pixels in the functional and spectral spaces, respectively (Carmona *et al*., 2016). However, even though addressing dissimilarities between communities functional and spectral structures is a more complete approach, addressing biodiversity using quantifiable and specific aspects of diversity (such as richness) is a widely used in research and it is fundamental in the understanding of many ecological processes (e.g., Carmona *et al*., 2021a; Swenson *et al*., 2016). Therefore, we estimated the relationships among log-transformed α–SD and both α–FD and α–TD using major axis regression within each transect (Legendre & Legendre, 2012). Additionally, we used the variation partitioning approach (Borcard *et al*., 1992; Legendre, 2008) to evaluate the unique and shared contribution of α–TD and α–FD in explaining α–SD variation. Variation partitioning was performed using the *varpart* function of the R package *‘vegan’* (Oksanen *et al*., 2020).

To explore in more detail the correspondence between functional and spectral spaces, we compared patterns in space occupation. Firstly, we visually compared functional and spectral structures of single transects by plotting their TPD functions at different probability quantiles. The value of a TPD function in each point of the space reflects the abundance of the corresponding combination of traits (in the functional space) or band values (in the spectral space). We graphically represented transects’ TPD for both FD and SD and highlighted contours containing 50, 75, and 99% of the total probability. Dissimilarities in transects’ functional and spectral structures were visually highlighted by plotting pairwise (functional and spectral) TPD functions. Later, we checked the correspondence between the functional and spectral structures by splitting herbaceous communities from the ones where a woody species (*Juniperus macrocarpa*) was dominant (≥ 50% relative abundance). Given that woody and herbaceous species occupy distinct portions of the global plant functional space (Díaz *et al*., 2016; Carmona *et al*., 2021a), if SD is detecting FD patterns, we expect to retrieve a similar a similar distinction in the pattern of occupation of the spectral space. Thus, we computed TPD functions for plots dominated by woody and herbaceous species in both functional and spectral spaces. Then, we plotted dissimilarities between woody and herbaceous functional and spectral structures.

### 2.6 Diversity patterns along the environmental gradient

In order to test whether taxonomical, functional, and spectral diversity patterns are coordinated along the sea-inland gradient, we performed a series of models along transects. First, we related all β–diversities to the interaction between spatial distance (i.e., Euclidean distance) and the type of diversity facet (i.e. TD, FD or SD). We computed spatial distance by progressively enumerating plots within transects according to the sea-inland gradient. Distances were normalized on their range between 0-1, with values closer to 0 indicating smaller distances between plots and vice versa. Considering the non-linear patterns occurring in coastal dune diversity along the sea-inland gradient (Acosta *et al*.,2007), we performed a Generalized Additive Model (GAM) for each transect using diversity facet as parametric term and spatial distance as the smooth term. The latter was computed using 5 basis functions (k) of thin plate regression splines basis type, specifying diversity facet as factor by variable. All GAMs were performed using REML method for the estimation of the smoothing parameter.

Then, we compared patterns of α–TD, α–FD, and α–SD along each transect. We standardized (zero mean and unit variance) α–diversity values for each diversity facet to have comparable ranges of variation. Then, we estimated relationships between α– diversity (response variable), and the interaction between distance from the sea and the type of diversity facet. We performed a GAM model per transect as specified above. All mentioned analyses were performed using R version 4.0.3 (R Core Team, 2020).

### 2.7 Partition of diversity across spatial scales

To test our hypothesis that the partition of SD across spatial scales would mirror that of FD, we partitioned species, functional and spectral richness using additive partition of diversity (Crist *et al*., 2003; Silvestre *et al*., 2021). We performed diversity partition considering the spatial organization of the sampling: a) α-diversity within plots (i.e., mean α_plot_), b);*β*-diversity between plots (i.e., α_transect_ – mean α_plot_), and c) *β*-diversity between transects (i.e., γ-diversity – α_transect_). We divided diversity values by their maximum to express the metrics in a range from 0 to 1.

## 3. Results

We recorded a total of 75 plant species in the study area (10.2 ± 2.8 std. dev. species per plot; Table S1). *Helichrysum* stoechas was the most abundant species accounting for ca. 24.2% of the total vegetation coverage, followed by the shrub *Juniperus macrocarpa* covering ca. 21.8% of total vegetation, and by Lomelosia rutifolia (ca. 11.2%).

The first principal component of the functional space (PC1; 51.76% of total variance) was mainly described by Ψ_tlp_, and LDMC, reflecting a trade-off between drought resistance and resource use, whereas PC2 (25.02% of total variance) was mainly related to venation architecture (i.e., VLA_minor_, Fig. 1).

**Fig. 1.**
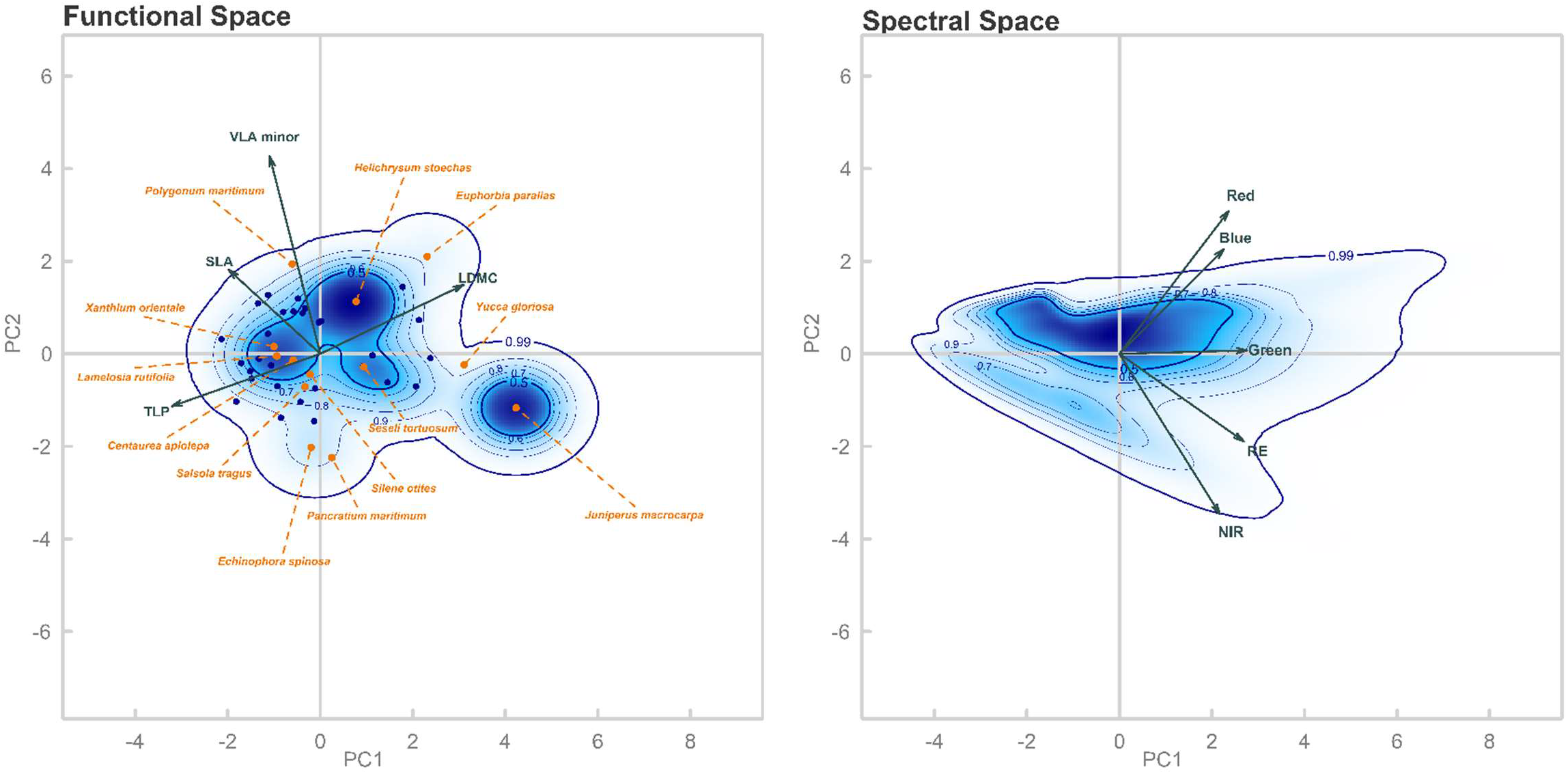
Probabilistic distribution of species in the functional space (left panel) and pixels in the spectral space (right panel) built using the first two PCA axis. Arrows lengths are proportional to the loadings of the considered traits and bands, respectively. Contour lines in each panel represent the thresholds of the probability density distributions (i.e., solid thin lines are 60%, 70%, 80%, 90%; bold lines are 50% and 99% of the total probability distribution). Probability distributions are based on the species and pixel present in the whole study area. The colour gradient highlights different probability densities with dark blue corresponding to portions of the space displaying the highest probabilities. Points present in the functional space represent species positions; a subset of species is highlighted in orange.

Regarding the spectral space, the first principal component was positively associated with the value of the green band; whereas blue and red bands had positive loadings on PC2 and were negatively associated with NIR and RE (Fig. 1). Both the functional and the spectral spaces displayed two main clusters of high probability density which, in the case of the functional space, clearly separated between woody (*Juniperus macrocarpa*) and herbaceous species.

### 3.1 Correspondence among functional structure, spectral structure, and species composition

β–TD ranged between 0 and 1 with a mean for all transects of 0.6 ± 0.19 (mean ± std. dev.) β–FD ranged between 0.02 and 0.99 (0.54 ± 0.21), β–SD varied between 0.07 and 1.00 (0.68 ± 0.22). We found significant positive correlations between dissimilarities of all diversity facets. The Mantel tests considering all plots of the study area showed a positive correlation between β–TD and β–SD (ρ = 0.20 p = 0.001). However, when controlling for β–FD, partial Mantel test was not statistically significant (ρ = 0.03, p = 0.088). The Mantel test between β–FD and β–SD revealed a positive correlation (ρ = 0.39, p = 0.001), which was confirmed also when controlling for β–TD (partial Mantel test ρ = 0.34, p = 0.001).

Considering dissimilarities within transects, all Mantel tests considering both β–TD (Fig. S1) and β–FD were significantly and positively correlated with β–SD as stressed by the major axis regression lines (Fig. 2). Both β–SD and β–FD were driven almost exclusively by high nestedness (i.e., high overlap) between plot structures (Fig. S2), suggesting that differences between plots from the same transect were mainly related to differences in the way species and pixels occupy the same areas of the functional and spectral spaces, respectively.

**Fig. 2.**
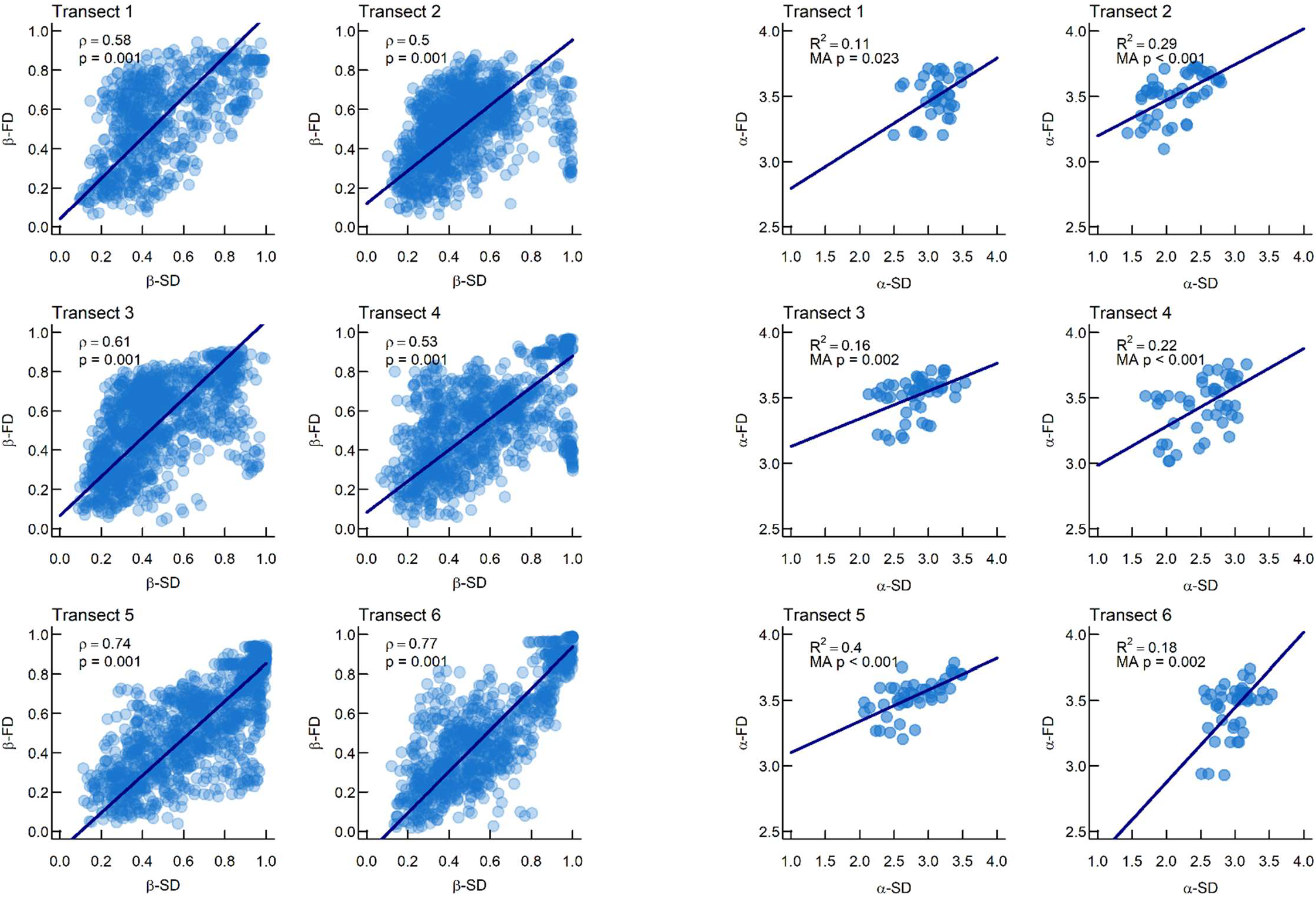
Relationships between transects’ SD and FD for both α-and β- diversity. The first two columns show β-diversity relationships, including the coefficient of correlation (ρ) and its significance (p; based on Mantel tests). Third and fourth columns show α–diversity relationships, including R^2^ and p-value (p) of the major axis regressions. In all plots, major axis regression line is showed in dark blue.

All major axis regression models relating α–FD to α–SD (Fig. 2) showed statistically significant positive relations, whose strength (R^2^ between 0.11 – 0.40) and slope were variable between transects. In contrast, all major axis regression models relating α–TD with α–SD within single transects failed to show significant relations with the only exception of transect 5 (Fig. S1). Variance partitioning of α–SD showed that most of its variation was accounted for variation in α–FD alone (10%). Joint effect of α–FD and α–TD explained almost no effect (1%), while α–TD did not explain any variation in α–SD.

Functional structures of transects showed consistent shapes and density distributions, suggesting that all transects shared similar combinations of traits (Fig. S3). Accordingly, functional dissimilarities between transects were low (0.2 ± 0.09 std. dev.; Fig. S4). The regions with the highest probability density (i.e., 50 % and 75% of the total functional structure) were consistent in all the transects, showing highly nested functional structures (nestedness approaching 1 in all pairwise comparisons, Fig. S4). Compared to functional structures, spectral structures showed higher variation among transects but highly consistent patterns were present in the regions with the highest probability density (i.e., 50 % and 75% of the total spectral structure; Fig. S3). The high consistency observed in the patterns reflecting high probability areas, was further corroborated by the low values of dissimilarity found between transects (0.48 ± 0.19 std. dev.; Fig. S4). The high values of nestedness found between transects (range 0.74 - 1; Fig. S4), confirmed that transect dissimilarities derived from different abundance of spectral characteristics that are common to pairs of transects, rather than by transects occupying non-overlapping areas of the spectral space (Fig. S4).

Comparisons of the functional and spectral structures of herbaceous-dominated and woody-dominated plots revealed high coordination between patterns in space occupation at the plot level (Fig. 3). For example, the high probability density clusters (i.e., the parts of the space that contain between 50% and 70% of the total probability) of woody and herbaceous plots occupied distinct and almost unique portions of both functional and spectral spaces. We found high dissimilarity values between woody and herbaceous plot structures (β–FD = 0.83, β–SD = 0.84), stressing the distinction between the set of traits and bands of woody and herbaceous communities (Fig. 3).

**Fig. 3.**
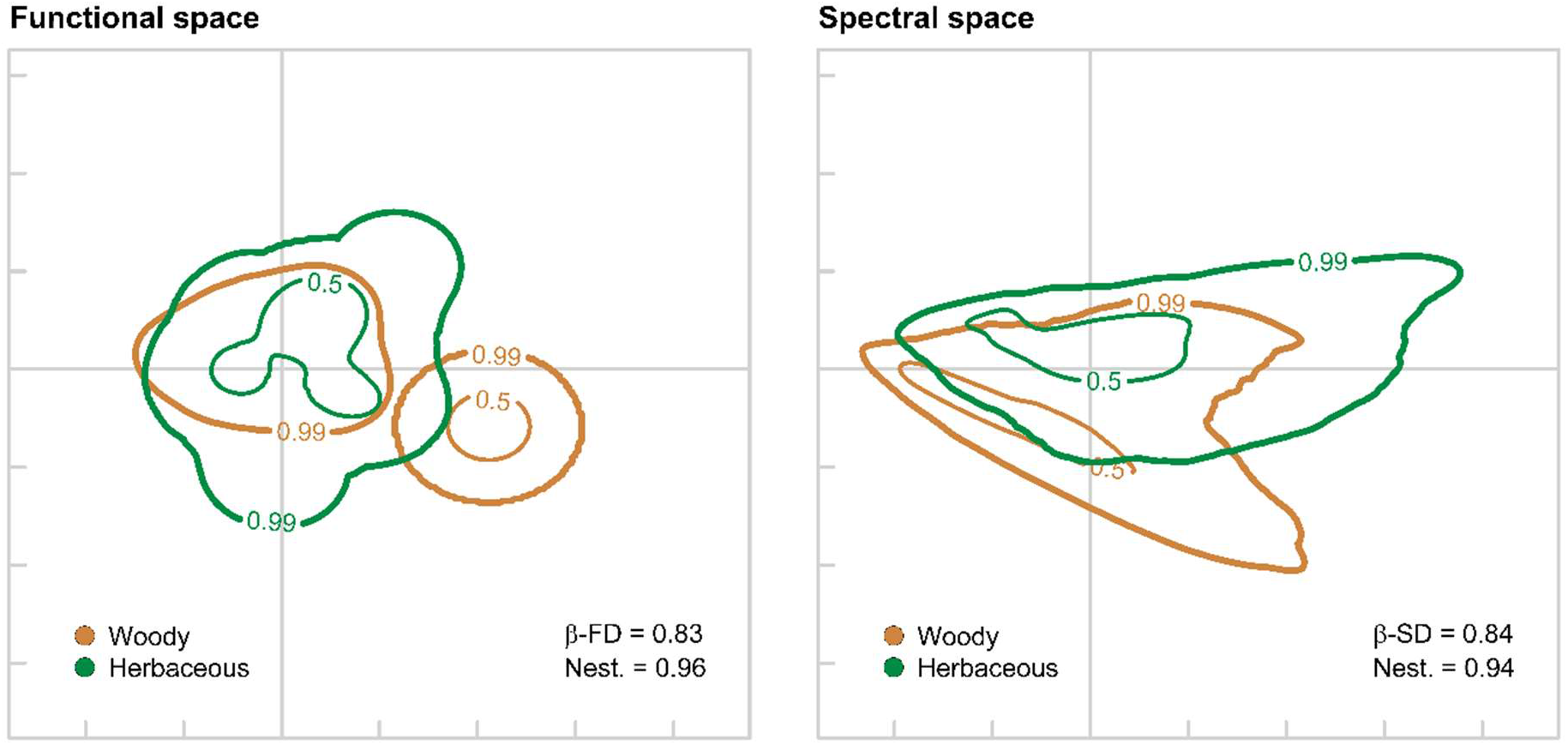
Probabilistic distribution of functional (i.e., FD, left panel) and spectral (i.e., SD, right panel) structures for *Juniperus*-dominated (Woody; brown lines) and herbaceous-dominated (Herbaceous; green lines) plots in functional and spectral spaces, showing the 50% and 99% quantiles of the TPD functions. The dissimilarity between woody and herbaceous plots was estimated using probabilistic overlap between their TPD functions in the functional and spectral spaces (β–FD and β–SD, respectively). Nestedness values (Nest.) represent the proportion of the dissimilarity that is due to differences in the density of occupation in the parts of the functional space that are shared between the woody and herbaceous TPD functions.

### 3.2 Diversity patterns along the environmental gradient

All analysed β–diversities relationship with the sea-inland gradient were statistically significant (p < 0.001; Fig. 4, Table S3). β–TD showed a generally a linear increase at increasing plot distances. β–FD showed a general increase at increasing plot distance which tend to flatten at higher values of functional dissimilarities in the majority of transects. β–SD showed transect dependent patterns, with a general increase at increasing plot distances.

**Fig. 4.**
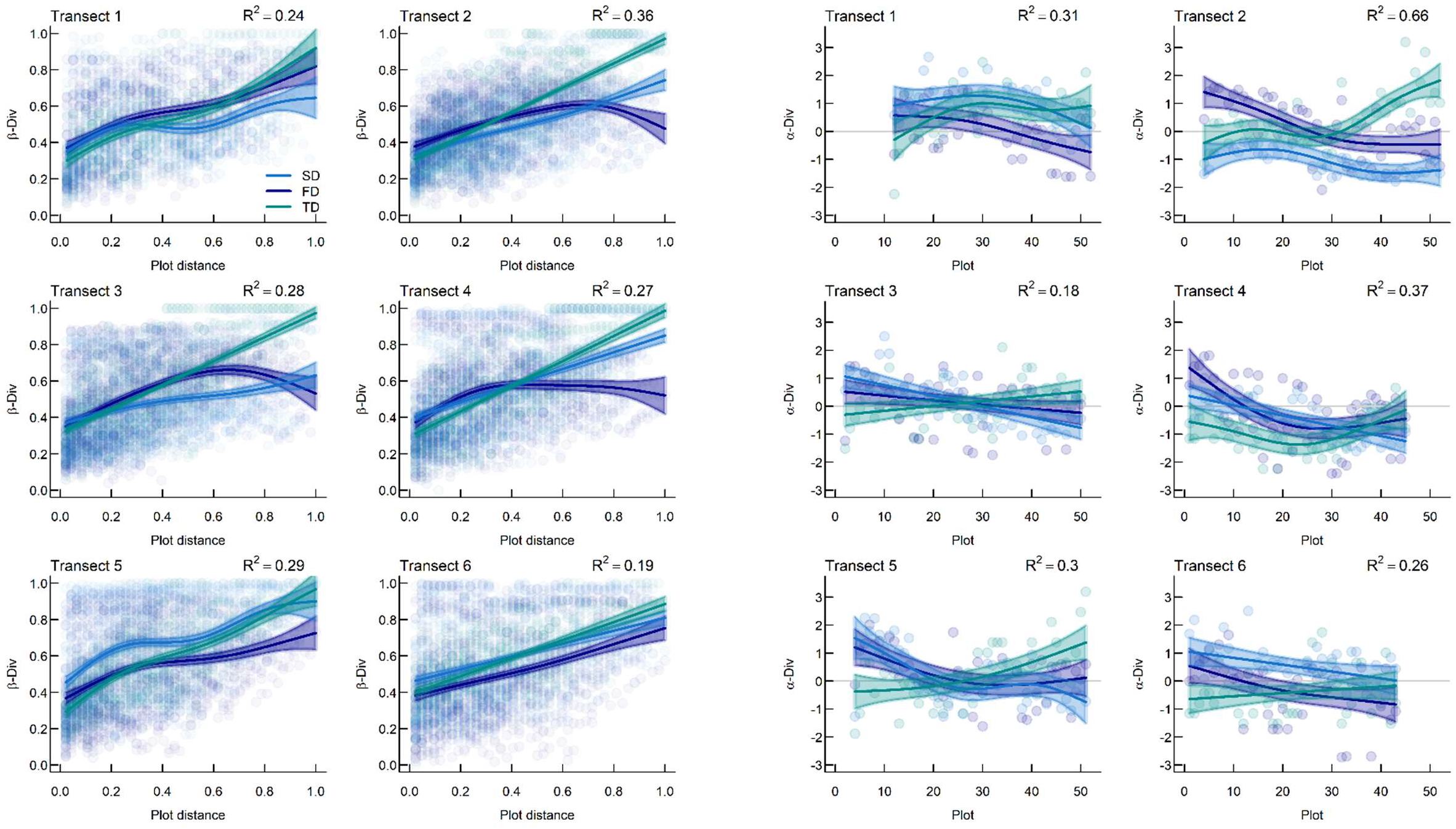
First two columns show the relationships between β–TD, β–FD, and β–SD and plot distances. Third and fourth columns show the patterns of α– TD, α–FD and α–SD along the sea-inland gradient. Plots are numbered increasingly from 1 (i.e., first plot in foredune) to a maximum of 52 (i.e., last plot in fixed dune) for each transect. All panels include fit line and R^2^ from GAMs; shaded area corresponds to 95% confidence interval for each diversity facet (SD: blue, FD: purple; TD: green).

α–diversity facets showed mainly significant relationships with the sea-inland gradient (Fig. 4, Table S2). α–TD showed a general positive non-linear trend along the sea-inland gradient, whereas α–FD showed a consistent slightly decreasing trend across transects. Likewise, α–SD mirrored α–FD along the sea-inland gradient, even though with variable extent depending on the transect under consideration.

### 3.3 Partition of diversity across spatial scales

Diversity facets partitioned differently across scales (Fig. 5). The highest portion of TD variation was found within and between transects (ca. 44.6 % and 42.2% of its total variation; respectively). Conversely, FD varied mostly within transect and especially at plot scale (ca. 60.5%). SD mirrored TD patterns, displaying the highest variation at transect level (i.e., β-diversity between plots and transects) accounting for about 77% of the total variation.

**Fig. 5.**
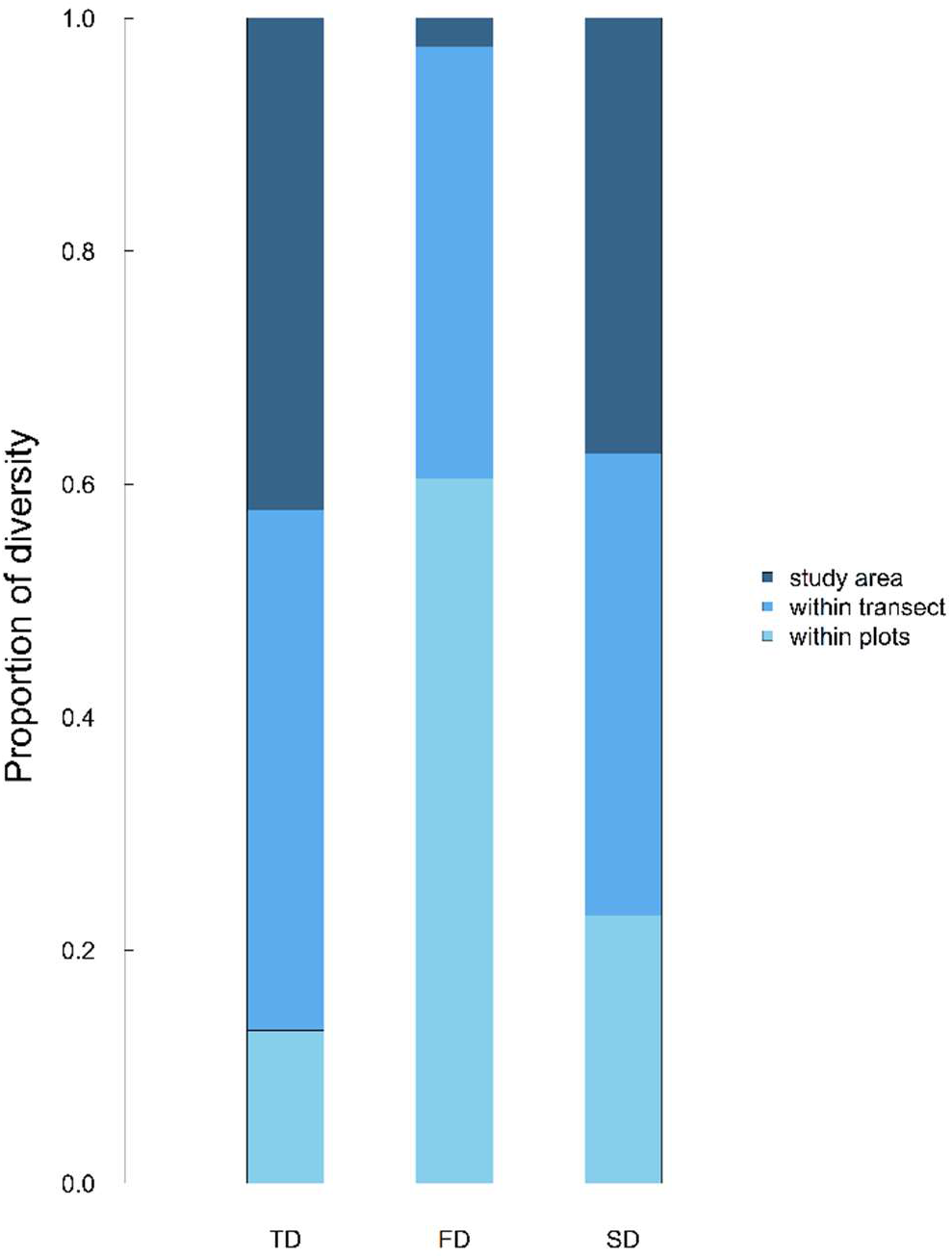
Taxonomical (TD), functional (FD), and spectral (SD) diversity partitions across spatial scales. Each column represents the total proportion of diversity detected in the whole study area (γ–diversity), divided in three different levels: “within plots” showing the proportion of diversity at plot level (α–diversity; azure colour); “within transect” showing the proportion of diversity at transect level (β-diversity between plots; light blue colour); and “study area” showing the proportion of diversity between transects (β-diversity between transects; dark blue colour).

## 4. Discussion

Approaches based on remote sensing data allows for affordable and continuous estimations of biodiversity across large areas, which is fundamental to cope with ongoing global change (Jetz *et al*., 2016; Reddy *et al*., 2021). Using fine-scale airborne multispectral remote-sensed images on coastal dune ecosystems, we explored the ability of remote sensed imagery in approximating diversity patterns across different spatial scale and along environmental gradient. For this, we expressed SD as a probabilistic distribution, effectively using a common analytical framework for both SD and FD that allows to seamlessly estimate changes in diversity across spatial scales while accounting for all components of diversity. We showed that SD was more strongly related to FD than to TD. We displayed a clear correspondence between the functional and spectral structure of plots and transects, although the partitioning of SD seemed to mirror the taxonomical one. These results suggest that fine-resolution SD is a good proxy of different facets of diversity, confirming its potential to detect spatial changes in diversity patterns.

### 4.1 Correspondence among functional structure, spectral structure, and species composition

As hypothesised, we found clear relationships between SD and FD both within- (α) and between-plot (β) diversity (Fig. 2). The strong correlations between functional and spectral dissimilarities imply that there is a strong correspondence between functional and spectral structures: changes in traits of communities corresponded to changes in band values. Results were consistent also considering univariate components of functional and spectral variation such as richness, although relationships were more variable in strength and slope depending on the transect under consideration (Fig. 2). Richness has a peculiar behaviour compared to full functional and spectral structures. This is because richness considers only the amount of space comprised in the 99% probability threshold and not how species (and pixels) occupy functional (and spectral) space (Carmona *et al*., 2016). Consequently, richness is strongly influenced by species (and pixels) composition, thus is less stable than the full structures of a community and could potentially be more influenced by the sampling resolution of a study (Pakeman, 2014; Carmona *et al*., 2016). Therefore, we hypothesize that the transect-dependant variability of functional and spectral relation may result from the resolution mismatch of functional and spectral samplings in our study.

Collecting functional traits can be a time-consuming process (Homolová *et al*., 2013) constrained by operative limits given by the number of species to sample, the replicates per individuals, the number and type of traits chosen, which all together contribute to define the functional space under investigation. In contrast, the nature of remote sensing samplings allows for the detection of the full set of traits influencing the spectral signal of each individual of the spectral image, depending on the spatial and spectral resolutions at which images are taken (Wang & Gamon, 2019; Cavender-Bares *et al*.,2020). Consequently, variation in SD derive from species presence, abundances, and interspecific variability of the complete set of features defining single individuals’ spectra. In our study, operative limits constrained the functional sampling to a species-level generalization of individuals, without including intraspecific trait variability. Conversely, SD was a snapshot of underlining individuals, with multiple replicates (i.e., pixels) per individual depending on individuals’ size. Therefore, for instance, plots containing the same species but occupying distinct positions in the sea-inland gradient might possess inter-individual differences in space occupation that spectral structure can account for, whilst the same level of detail cannot be reached by the functional structure as estimated in this study (but see Wong & Carmona, 2021 for alternative approaches to incorporate intraspecific trait variation in FD estimations). As a result, this mismatch in sampling resolution may influence the strength of the relationship, producing different R^2^ across transects.

However, despite the difference in resolution between functional and spectral samplings in this study, the strong relation between SD and FD suggests that SD is indeed able to retrieve the functional structure of the studied communities. Consistently, functional and spectral structures of woody- and herbaceous-dominated plots showed strong correspondence, demonstrating that differences in functional space occupation between woody and herbaceous species (as also shown in the global spectrum of plant form and function; Carmona *et al*.,2021a; Díaz *et al*.,2016) are analogous also in the correspondent spectral space (Fig. 3). This similarity in space occupation stress the potential of SD in approximating multivariate FD patterns, resulting promising to directly assess community and ecosystem changes (Díaz *et al*.,2016; Joswig *et al*.,2022). Consistent results were found also across spatial scales, with transect spectral structures mirroring the same pattern of the functional ones (Fig. S3-S4).

The lack of a significant relationship between α–TD and α–SD as well as the influence of β–FD in defining β–TD and β–SD correlation, suggested that fine resolution multispectral SD failed to detect the taxonomical component of biodiversity (Fig. S1). This result contrasts with previous research on the same ecosystems (see Marzialetti *et al*.,2021 which used a 3m pixel resolution), but can be explained by the resolution of the images used in our study. The fine pixel resolution we used (~ 3 cm) provides leaf details which might better approximate leaf-level functions (Cavender-Bares *et al*.,2017), while failing in capturing the features that additively define a species as a taxonomic entity. Whereas for the estimation of species richness all species are completely unique, this is not the case for functional or spectral diversity, where we need to consider the role of redundancy (Cadotte *et al*.,2011). Different species can occupy similar positions in the functional/spectral space, irrespective of their taxonomic identities, influencing TD-SD relationships (Schmidtlein & Fassnacht, 2017; Fassnacht *et al*.,2022). Consistently, β–diversity correlations lacked a clear and significant link between the spectral and taxonomical component, pointing toward a hidden influence of FD as highlighted by partial Mantel tests and variation partitioning. However, we cannot exclude that finer spectral resolution (i.e., hyperspectral) could be capable to discriminate the whole set of traits needed to identify the spectral signal of a species as a distinct taxonomic unit (Ustin *et al*., 2009; Gamon *et al*., 2020).

### 4.2 Diversity patterns along the environmental gradient

Patterns along the sea-inland gradient were in agreement with what already described in literature (Acosta *et al*., 2008), both in its α- and β- taxonomical component.

The saturation of higher plot functional dissimilarities at increasing plot distances (Fig. 4), suggests that the higher portion of β–FD variation is independent from the distance between plots. Indeed, species co-occurring in foredune plots (e.g. *Xanthium orientale, Euphorbia paralias, Echinophora spinosa*) as well as species co-occurring in inland plots (e.g. *Juniperus macrocarpa, Seseli tortuosum, Helichrysum stoechas*; Ciccarelli & Bona, 2022) show a different set of traits, suggesting a differentiation in functional strategies of co-occurring species (Fig. 1). Therefore, depending on the species composition, adjacent plots may possess higher dissimilarities than distant ones, leading to weak relationships with the sea-inland gradient. However, in agreement with Carboni *et al*. (2013), α–FD decreased proceeding landward, suggesting that species closer to the sea are more functionally different than inland ones. Indeed, foredune communities are less redundant in species functional strategies (e.g. *Pancratium maritimum, Salsola tragus, Xanthium orientale*) compared to inland ones (Silene otites, Centaurea aplolepa subsp. subciliata, Hieracium picenorum; Acosta *et al*., 2006; Fig.1), suggesting that plants can cope with harsh environmental conditions through different ecological strategies (Rota *et al*., 2017).

Depending on the considered transect, α–SD along the sea-inland gradient mirrored FD, whereas β–diversity relationships showed a more linear increase than the functional one (Fig. 4). As discussed above, this mismatch between functional and spectral variation may result from the resolution mismatch between functional and spectral samplings. Moreover, we should consider that plant spectral variation depends on vegetation size (Conti *et al*., 2021) and complexity (Hauser *et al*., 2021) so that larger individuals occupy larger portions of the spectral space (Schweiger *et al*., 2021). Therefore, differences in individuals’ size, vertical complexity and traits along the sea-inland gradient (Tordoni *et al*., 2019) may increase SD variation, amplifying dissimilarity between plots sharing the same species in a way that cannot be mirrored by the FD sampling as performed here. Altogether, these differences result in slightly different patterns of SD and FD along the sea-inland gradient.

### 4.3 Partition of diversity across spatial scales

In agreement with previous studies (Del Vecchio *et al*.,2018; Tordoni *et al*.,2018), we observed that the highest portion of taxonomical variation occurred at transect level (Fig. 5). This can be explained by differences in dune structure that produce small-scale differences in abiotic conditions along the sea-inland gradient which, in turns, generate the observed differences in species richness at broader scales (Acosta *et al*.,2008; Tordoni *et al*.,2018).

In contrast, more than 60% of the FD variation was found at plot level (Fig. 5), showing that differences in traits between co-occurring species are comparable to differences found at broader spatial scales. This taxonomical-functional disequilibrium in diversity partition was already observed in other ecosystems (de Bello *et al*.,2009; Carmona *et al*.,2012) and across taxonomical groups (Silvestre *et al*.,2021). Accordingly, we detected consistently higher functional differences between plots than between transects (Fig. S2 – S4). The prominent role of nestedness in explaining plots’ functional dissimilarities further corroborated this pattern: indicating that functional differences between pairs of plots were due to differences in how densely the same areas of the functional space where occupied, rather than each plot occupying exclusive areas of the functional space. Transects showed similar patterns in functional space occupation, further stressing that most of the realisable trait combinations of dune ecosystems were already expressed at smaller spatial scales. This result is in line with recent efforts showing that FD is highly preserved within ecosystems even at the global scale (e.g. Testolin *et al*.,2021), stressing the effect of lower scale abiotic filters in the selection of the suite of traits occurring in a given site.

Compared to FD, we found larger proportions of SD variation at broader scales (i.e., among plots and transects, Fig. 5). Surprisingly, we also detected a difference in partitioning between FD and SD contrarily to our expectations, but this may be reconciled by what already explained above about sampling mismatch between these facets. Indeed, being SD the product of multiple within-individual replicates, it could potentially exacerbate inter- and intra-individual variation that will magnify lower spatial scale differences, while additively contributing to define broader scales species’ spectra. Higher β–SD were observed at plot level compared to transect ones, confirming that spectral variation is realized within transects rather than between them. Additionally, spectral dissimilarities between pairs of transects were mostly due to differences in the areas of the spectral space that were not very densely occupied, whereas the high density portions (spectral “hotspots”*sensu* Carmona *et al*.,2021b) showed a high degree of overlap (Fig. S3 - S4) suggesting that transect variability results mostly from differences in the expression of the same spectral features.

In other words, vegetation from different transects shares a recurrent set of functional traits that can be remotely detected by SD.

## 5. Conclusion

Biodiversity detection through remote sensing is a promising tool for addressing ongoing global changes. Yet, before we can consider this tool as a reliable one for biodiversity monitoring, major gaps on the relationships between biodiversity facets and the remote sensed signal should be filled. Here, we applied to spectral diversity an approach (TPD) that allows examining all aspects of functional diversity for multiple traits by applying a probabilistic formalization. We show that using the TPD approach for estimations of spectral diversity allows to consider all aspects of spectral variation simultaneously, so that both FD and SD can be analysed with a congruent set of analytical methods. Using these methods, we showed that SD consistently covaried with FD at different spatial scales. By contrast, we did not observe a similar degree of covariation between SD and TD, suggesting that detecting the taxonomical signal is a more complex task, probably due to many species being redundant in their spectral signal. Despite the ability of SD to detect FD patterns, we found that while most of FD variation was found at the plot level, SD variation was more homogeneously distributed across spatial scales, probably because the spectral signal incorporates intra- and inter-individual differences. Since this same level of sampling detail is not always achievable when collecting functional trait data, SD appears as a powerful surrogate to estimate the functional structure of plant communities while accounting for the relative contribution of intraspecific trait variability.

## Acknowledgments

CPC and EB were supported by the Estonian Research Council grant (PSG293) and the European Regional Development Fund via the Mobilitas Pluss programme (MOBERC40). ET is supported by Estonian Research Council grant MOBJD1030. We would like to thank Dr. Francesca Logli along with the Migliarino-San Rossore-Massaciucoli Regional Park administration that authorized the field sampling activities. We are extremely grateful to Andrea Nardini that allowed us to use his laboratory and equipment to measure functional traits. We also thank Erika Bellini for her help in handling sampling materials.

## Author contributions

G.B., C.P.C., E.T., and F.P. conceived the study. E.B., E.T., F.P., G.B., and N.P. collected plant and functional trait data. D.M. and G.C. collected spectral data. E.B. and N.P. measured functional traits. E.B., D.M., and G.C. pre-processed spectral data. E.B., E.T., C.P.C. performed the analysis. E.B., E.T., C.P.C., F.P., D.R., M.D., D.C. and G.B. interpreted the data. E.B. wrote the first draft of the manuscript and all authors substantially contributed to revisions.

## Data availability statement

In case of paper acceptance, the data needed to reproduce the main results will be made available in a public repository (e.g., figshare, zenodo).

