## Supplementary information for "Plant spectral diversity from high-resolution multispectral imagery detects functional diversity patterns in coastal dune communities"

### Supplementary methods

*Functional traits measurements*

We measured leaf functional traits associated with the "Leaf Economics Spectrum" (LES;(Wright *et al.*, 2004), which reflects the trade-offs between potential photosynthetic carbon gain and carbon investment for leaf construction. We also measured hydraulic traits strictly related to water transport efficiency (leaf venation architecture, (Sack & Scoffoni, 2013) and drought resistance (osmotic potential at full turgor and water potential at turgor loss point, (Bartlett *et al.*, 2012), which are coordinated in the hydraulic safety-efficiency trade-off (see Petruzzellis *et al.* 2021). Following Petruzzellis *et al.* (2021), we selected the following functional traits: Specific Leaf Area (SLA, $mm^2/mg$); Leaf Dry Matter Content (LDMC, mg/g); Minor Vein Length per unit Area ($VLA_{minor}$, $mm/mm^2$); Water potential at turgor loss point ($\Psi_{tlp}$, - MPa).

Entire individuals were collected in the field, wrapped in cling film, placed in plastic bags with a humid paper inside, and stored in cool bags until their transport (within 3 hours) to the laboratory. Once in the laboratory, individuals were rehydrated overnight by placing the rooting apparatus in glass containers with tap water.

For the measurement of SLA and LDMC, leaves were randomly selected and detached from each individual, and the petiole was removed. For each individual, 2 to 15 leaves were sampled to reach a substantial mass to measure the dry weight. Leaf turgid weight was measured using an analytical balance, then each leaf was scanned, and total leaf area was calculated using Image J software (Rasband, 2018). Subsequently, leaves were oven dried at 70 °C for 48 h and their dry weight was then measured. Then SLA was calculated as Leaf Area/Leaf Dry Weight [$mm^2/mg$] and LDMC as Leaf Dry Weight/Leaf Turgid Weight [mg/g] (Pérez-Harguindeguy *et al.*, 2013).

$VLA_{min}$ was calculated as Vein Length/ Leaf Sample area [$mm/mm^2$]. One leaf was randomly selected from each individual, and samples were obtained by cutting a smaller part (c.a. 3 $cm^2$), avoiding the midrib. Samples were put in 1 M NaOH solution for 48-72 hours at room temperature to dissolve pigments. NaOH solution was carefully replaced when it turned from transparent to dark-colored. After that, samples were washed in distilled water and bleached in NaClO 5% for 15/30 minutes. Then, leaves were gently dried with a piece of blotting paper and leaf hairs were carefully removed. Samples were then treated in a sequence of ethanol solutions at an increasing concentration (25%, 50%, 70%, 100%) and then colored in an alcoholic solution of toluidine blue (3%) overnight. Samples were then washed in a series of ethanol solutions at decreasing concentrations to remove excess dye. Microscopic slides were prepared with samples of approximately 1.5 $cm^2$. Images were then captured using an optical microscope (4X magnification) equipped with a digital camera (model Syrio-2, PbInternational) and $VLA_{min}$ was measured using PhenoVein software (Bühler *et al.*, 2015).

Leaf osmotic potential at full turgor ($\pi_0$) and water potential at turgor loss point ($\Psi_{tlp}$) were measured according to the procedures described in Petruzzellis *et al.* (2021). Since $\Psi_{tlp}$ was calculated based on $\pi_0$, only $\Psi_{tlp}$ was considered for the analyses.

*Remotely sensed image acquisition and processing*

We conducted Unmanned Aerial Vehicle survey with a DJI Phantom 4 Pro v.2 (DJI, Shenzhen, China) equipped with a 1" CMOS RGB sensor of 20 Mpix and a MicaSense RedEdge 3© multispectral camera (MicaSense, Seattle, WA) with a 1280 × 960 global shutter sensor.

| ***Band number*** | ***Band name*** | ***Center Wavelenght (nm)*** | ***Bandwidth FWHM (nm)*** |
|---|---|---|---|
| 1 | Blue | 475 | 20 |
| 2 | Green | 560 | 20 |
| 3 | Red | 668 | 10 |
| 4 | Near IR | 840 | 40 |
| 5 | Red Edge | 717 | 10 |

To take pictures under similar light conditions, flights were conducted during two consecutive sunny days (29 – 30 May 2019) between 10:00 and 16:00. During each flight, we collected high-resolution visible (RGB) and multispectral photos. Flight plans were planned with more than 80% of sidelap and overlap to provide significant frame overlap for enhanced SfM (Structure from motion) processing. A total of 11.2 ha of terrain were surveyed at a flying altitude of 40 m AGL (Above Ground Level), 581 high resolution RGB images and 1295 high resolution multispectral images were taken. To obtain high precision in the georeferencing photogrammetrical model, an appropriate number of Ground Control Points (GCPs) had to be strategically placed. A Stonex S9III Plus NRTK-GNSS system connected to the Hexxagon Smartnet reference stations network for real-time positioning correction was used to measure each GCP. On GCPs, the accuracy (measured as Root Mean Square Error) was roughly 2 or 3 centimeters. To achieve high accuracy in conversion of altimetric value from ellipsoidical model to geoidical one, we based the conversion on the Datum IGM42, using GK2 IGMI (*Istituto Geografico Militare Italiano*) grids.Using Agisoft Metashape Professional, the captured images were first pre-processed (brightness correction, alignment, calibration) then analyzed to create a 3D model with high accuracy and resolution. From 3D model we generated:

- Orthophotos RGB with a ground sampling distance of about 1 centimeter for pixels.
- Orthophotos multispectral with a ground sampling distance of about 2 centimeters for pixels.

To remove all pixels not related to vegetation, we performed an unsupervised linear spectral unmixing process (Settle & Drake, 1993). Spectral unmixing (Shi & Wang, 2014) is a process that allows the recognition of image endmembers (i.e., pixels exclusively detecting only one type of underlying material such as vegetation) and calculates the percentage of each endmember present in each pixel. Spectral unmixing was performed on each multispectral image using Spectral Hourglass Wizard (SHW) application in ENVI software (v.3.4, Exelis Visual Information Solution, Boulder, Colorado). Spectral unmixing was performed on each multispectral image using Spectral Hourglass Wizard (SHW) application in ENVI (Environment for Visualizing Images) software (v.3.4, Exelis Visual Information Solution, Boulder, Colorado). SHW application guides the user step-by-step through the ENVI hourglass processing flow to find and map image spectral endmembers from hyperspectral data. For each multispectral image, the algorithm returns one image for each endmember, in which each pixel shows the fraction of the endmember material found in the image. These images were further used to build binary masks in QGIS 3.4 (QGIS Development Team 2020) that restrained only pixels containing at least 60% of the vegetation. Each complete multispectral spectral image was then cut using the previously produced binary mask, visually checked for errors and corrected where necessary. Pixels present in each plot of each transect were extracted using ‘raster’ (Hijmans, 2021) R package.

#### Supplementary material references:

**Bartlett MK, Scoffoni C, Sack L**. **2012**. The determinants of leaf turgor loss point and prediction of drought tolerance of species and biomes: a global meta-analysis. *Ecology Letters* **15**: 393–405.

**Bühler J, Jahnke S, Hülskamp M, Schurr U, Scharr H, Huber G, Rishmawi L, Pflugfelder D, Koornneef M**. **2015**. *phenoVein - A software tool for leaf vein segmentation and analysis*. Pflanzenwissenschaften.

**Hijmans RJ**. **2021**. *raster: Geographic Data Analysis and Modeling*.

**Pérez-Harguindeguy N, Díaz S, Garnier E, Lavorel S, Poorter H, Jaureguiberry P, Bret-Harte MS, Cornwell WK, Craine JM, Gurvich DE, *et al.*** **2013**. New handbook for standardised measurement of plant functional traits worldwide. *Australian Journal of Botany* **61**: 167.

**Petruzzellis F, Tordoni E, Tomasella M, Savi T, Tonet V, Palandrani C, Castello M, Nardini A, Bacaro G**. **2021**. Functional differentiation of invasive and native plants along a leaf efficiency/safety trade-off. *Environmental and Experimental Botany* **188**: 104518.

**Rasband WS**. **2018**. ImageJ. US National Institutes of Health, Bethesda, Maryland, USA. http://imagej. nih. gov/ij/.

**Sack L, Scoffoni C**. **2013**. Leaf venation: structure, function, development, evolution, ecology and applications in the past, present and future. *New Phytologist* **198**: 983–1000.

**Settle JJ, Drake NA**. **1993**. Linear mixing and the estimation of ground cover proportions. *International Journal of Remote Sensing* **14**: 1159–1177.

**Shi C, Wang L**. **2014**. Incorporating spatial information in spectral unmixing: A review. *Remote Sensing of Environment* **149**: 70–87.

**Wright IJ, Reich PB, Westoby M, Ackerly DD, Baruch Z, Bongers F, Cavender-Bares J, Chapin T, Cornelissen JHC, Diemer M, *et al.*** **2004**. The worldwide leaf economics spectrum. *Nature* **428**: 821–827.

### Figures

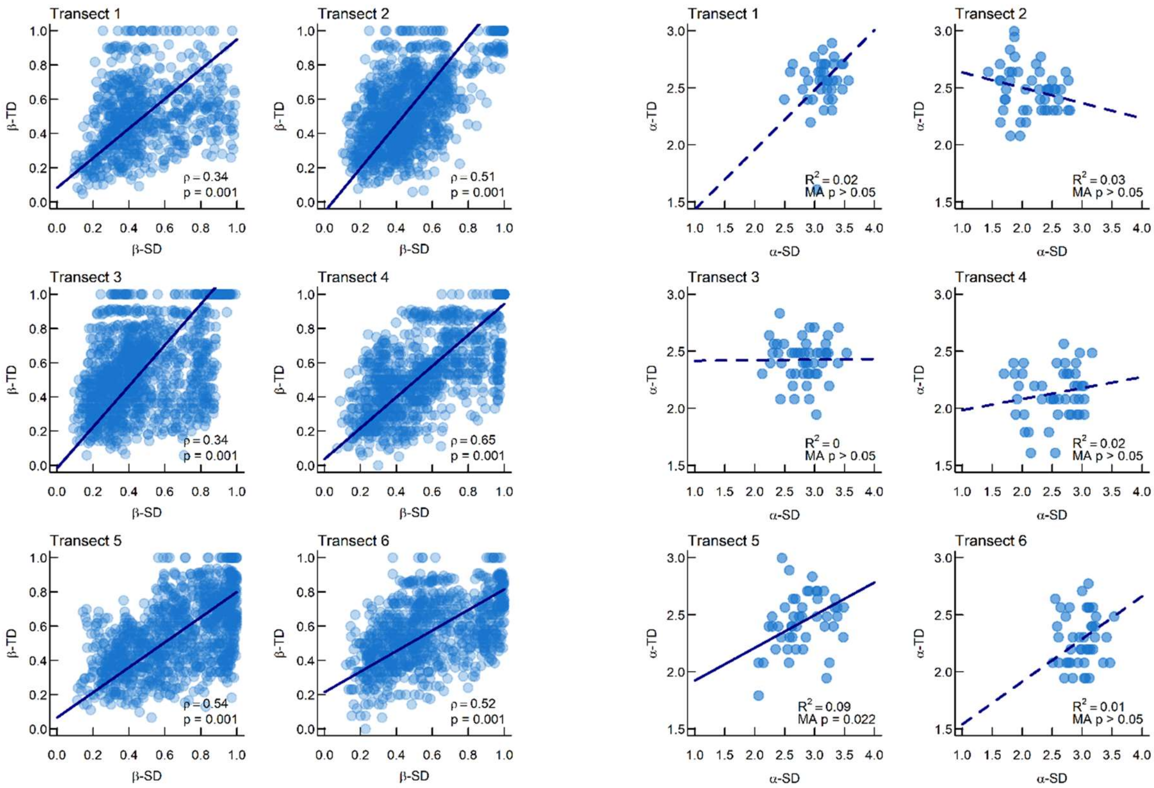

**Fig. S1** Relationship between transects' SD and TD for both α-and β- diversity. The first two columns show β–diversity relationships: Mantel test coefficient (ρ) is reported together with its significance (p) in the lower part of the graph. Third and fourth column show α–diversity relations: $R^2$ and p-value (p) of the Major axis regression are reported in lower part of the graph. In all plot, Major axis regression line is showed in dark blue: dashed lines are used for not significant regressions.

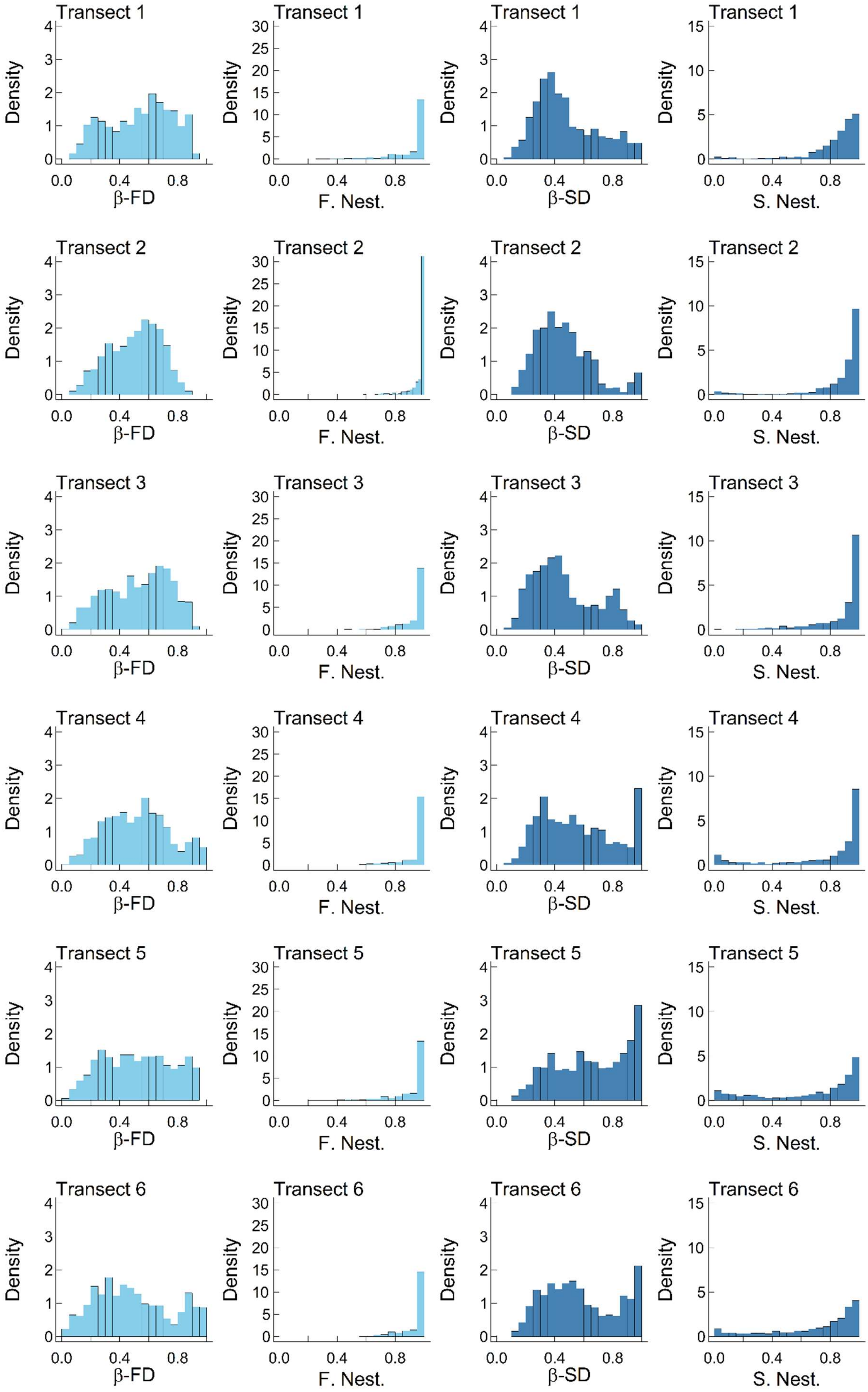
Transect 1
Transect 2
Transect 3
Transect 4
Transect 5
Transect 6
Density
β-FD
F. Nest.
β-SD
S. Nest.
0.0
0.4
0.8

**Fig. S2** Frequencies of β-FD (first column) and their related nestedness (Nest), β-SD (third column) values, and their nestedness (Nest) for each transect (rows from 1 to 6). Dark blue refers to spectral values, whereas light blue refers to functional values.

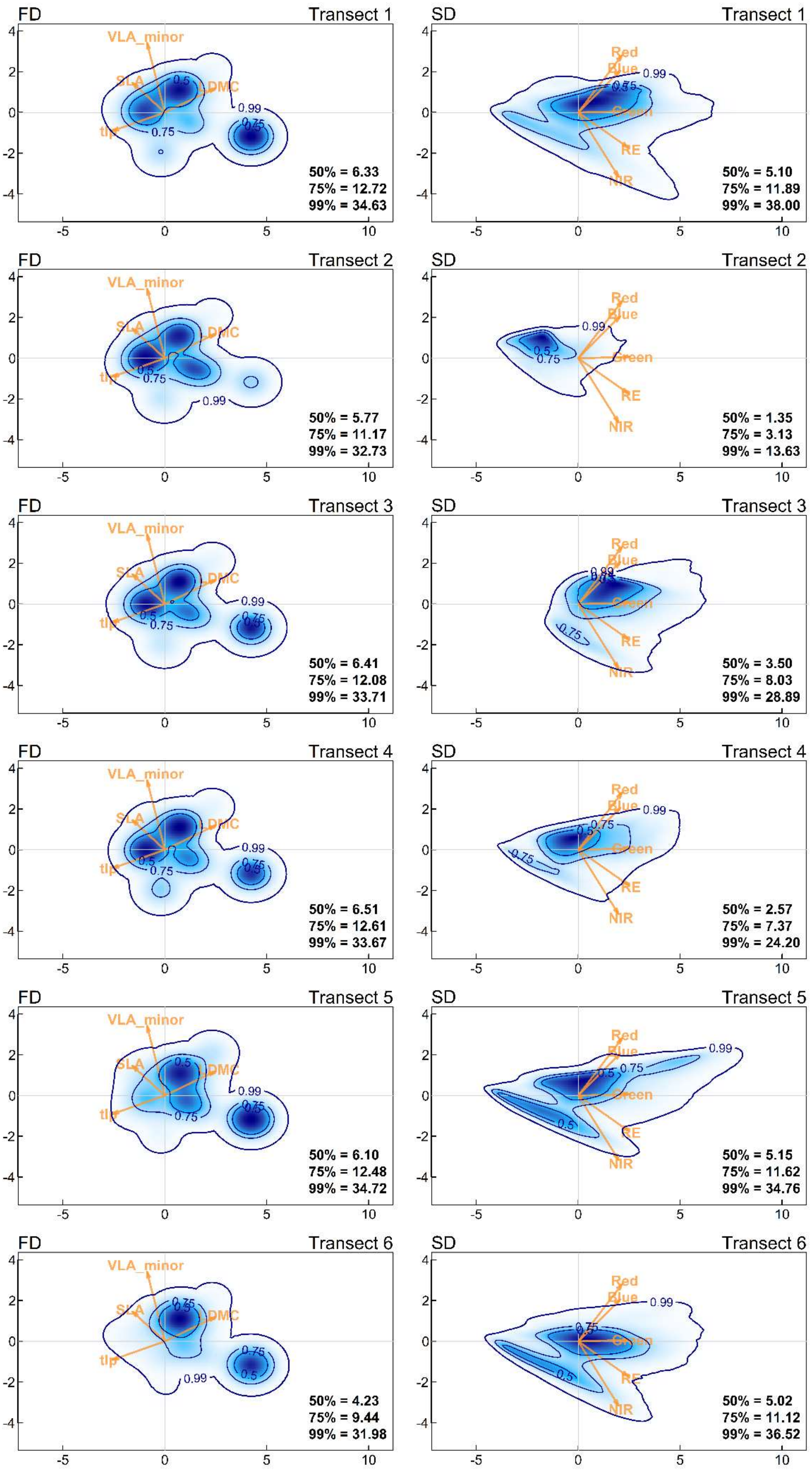
FD Transect 1
VLA_minor
SLA
DMC
tlp
50% = 6.33
75% = 12.72
99% = 34.63
SD Transect 1
Red
Blue
Green
RE
NIR
50% = 5.10
75% = 11.89
99% = 38.00
FD Transect 2
50% = 5.77
75% = 11.17
99% = 32.73
SD Transect 2
50% = 1.35
75% = 3.13
99% = 13.63
FD Transect 3
50% = 6.41
75% = 12.08
99% = 33.71
SD Transect 3
50% = 3.50
75% = 8.03
99% = 28.89
FD Transect 4
50% = 6.51
75% = 12.61
99% = 33.67
SD Transect 4
50% = 2.57
75% = 7.37
99% = 24.20
FD Transect 5
50% = 6.10
75% = 12.48
99% = 34.72
SD Transect 5
50% = 5.15
75% = 11.62
99% = 34.76
FD Transect 6
50% = 4.23
75% = 9.44
99% = 31.98
SD Transect 6
50% = 5.02
75% = 11.12
99% = 36.52

**Fig. S3** Probabilistic distribution of functional (i.e., FD, left panels) and spectral (i.e. SD, right panels) structures of each transect. Contour lines in each panel indicate the thresholds of the probability density distribution (i.e., 50%, 75% and 99% of the total probability distribution) representing the probability of finding underlying combination of traits (FD) or bands (SD). The legend of each panel contains the amount of space (i.e., functional and spectral richness, respectively) occupied for each probability threshold for each transect. The colour gradient highlights different probability densities with dark blue corresponding to the highest probabilities.

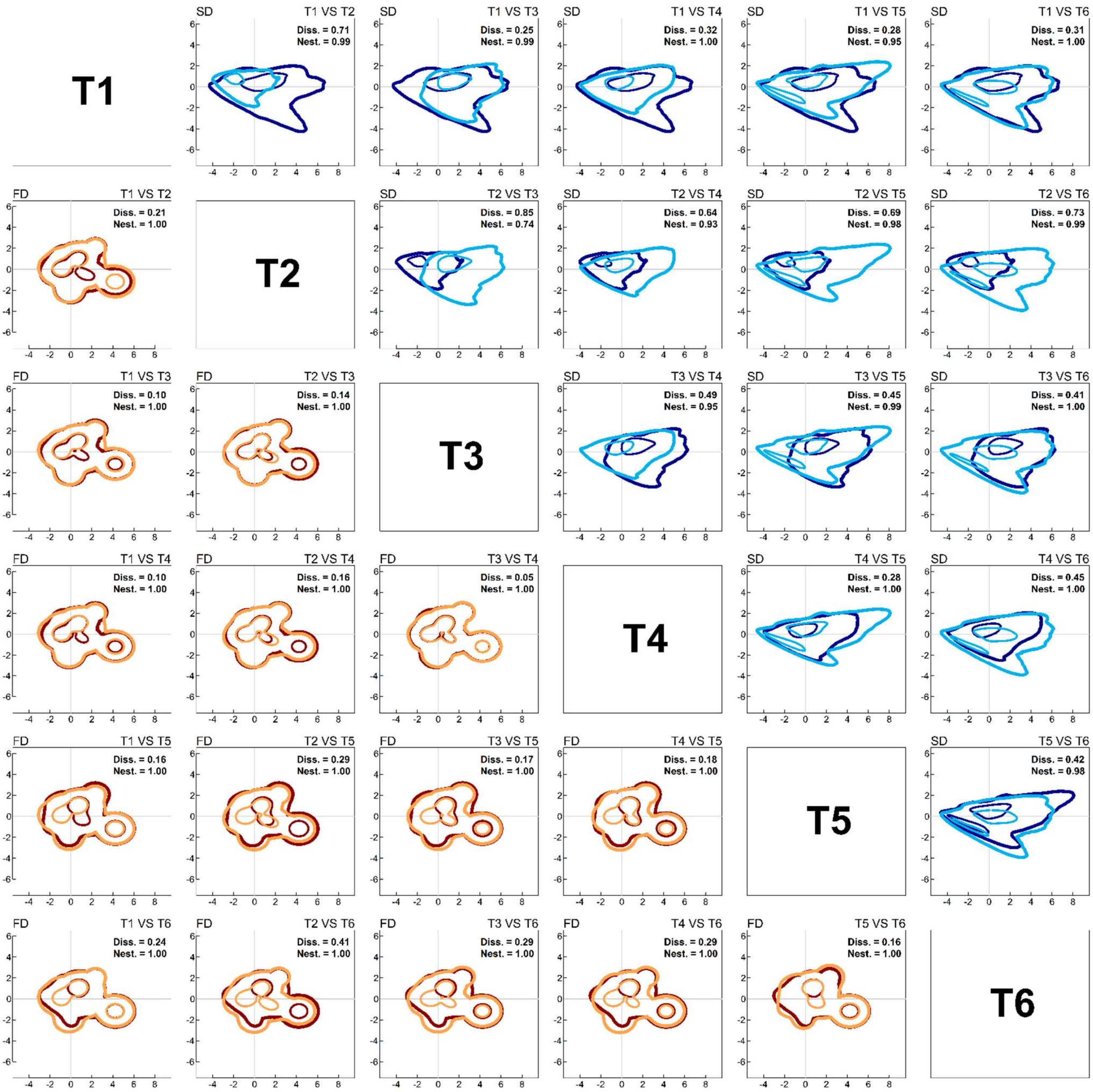

**Fig. S4** Pairwise overlap used to compute transect dissimilarity for both functional (lower triangle; red/orange colour palette) and spectral transect spaces (upper triangle; blue colour palette). Each probability density distribution is highlighted at 50% (inner thin line) and 99% (outer thick line) of total probability. Overlapping portions of the space represents the nestedness of the two transect spaces, highlighting combination of traits (for functional spaces) or band values (for spectral spaces) that are shared between pairwise transects. Both functional and spectral dissimilarities (Diss.) and nestedness (Nest.) values for each pairwise overlap are reported in the upper inset within each graph.

### Tables

**Table S1** Summary statistics (minimum, mean, maximum and standard deviation) for plots' α–TD, α–FD, and α–SD. For each diversity metrics minimum, maximum, mean, and standard deviation are shown for each transect and overall study area (Area).

| α–TD | **T1** | **T2** | **T3** | **T4** | **T5** | **T6** | **Area** |
|---|---|---|---|---|---|---|---|
| Min | 4.00 | 7.00 | 6.00 | 4.00 | 5.00 | 6.00 | 4.00 |
| Mean | 12.03 | 11.21 | 10.49 | 7.64 | 10.79 | 9.07 | 10.18 |
| Max | 17.00 | 19.00 | 16.00 | 12.00 | 19.00 | 15.00 | 19.00 |
| Std.Dev | 2.41 | 2.61 | 2.02 | 1.88 | 2.87 | 2.36 | 2.76 |
| α–FD | **T1** | **T2** | **T3** | **T4** | **T5** | **T6** | **Area** |
| Min | 23.63 | 21.14 | 22.96 | 19.43 | 23.66 | 17.72 | 17.72 |
| Mean | 32.58 | 32.89 | 32.94 | 30.86 | 33.48 | 30.71 | 32.27 |
| Max | 40.28 | 40.56 | 39.90 | 41.94 | 43.03 | 41.07 | 43.03 |
| Std.Dev | 5.13 | 5.30 | 4.64 | 6.33 | 4.54 | 5.60 | 5.34 |
| α–SD | **T1** | **T2** | **T3** | **T4** | **T5** | **T6** | **Area** |
| Min | 11.12 | 3.19 | 7.38 | 4.44 | 6.87 | 11.30 | 3.19 |
| Mean | 22.35 | 8.14 | 16.72 | 12.62 | 16.57 | 19.48 | 15.72 |
| Max | 34.33 | 15.38 | 33.23 | 22.75 | 31.56 | 33.31 | 34.33 |
| Std.Dev | 5.29 | 3.42 | 5.85 | 5.01 | 6.86 | 5.15 | 7.00 |

**Table S2.** Generalized additive model (GAM) results for all α–diversity relations with the sea-inland gradient. Each box contains a single model output for the considered transect. Parametric coefficients are reported in the upper part of the box together with their estimate, standard error and p-value. α– SD was set as reference category for each model. A separate smooth function was performed for each facet of α–diversity. Smooth terms are reported in the lower part of the graph together with their effective degree of freedom (edf) and p- values.

| | *Parametric coefficients* | | | *Transect* | *Parametric coefficients* | | | *Transect* | *Parametric coefficients* | | | *Transect* |
|---|---|---|---|---|---|---|---|---|---|---|---|---|
| | *Estimate* | *Std. Error* | *p-value* | | *Estimate* | *Std. Error* | *p-value* | | Estimate | Std. Error | p-value | |
| ***Intercept*** | 0.949 | 0.126 | 0.000 | | -1.083 | 0.091 | 0.000 | | 0.143 | 0.105 | 0.174 | |
| ***FD*** | -0.891 | 0.178 | 0.000 | | 1.198 | 0.128 | 0.000 | | -0.018 | 0.148 | 0.903 | |
| ***TD*** | -0.281 | 0.178 | 0.118 | | 1.455 | 0.128 | 0.000 | | -0.033 | 0.148 | 0.825 | |
| | ***Smooth terms*** | | | 1 | ***Smooth terms*** | | | 2 | ***Smooth terms*** | | | 3 |
| | *edf* | *p-value* | | | *edf* | *p-value* | | | *edf* | *p-value* | | |
| ***Plot:SD*** | 2.086 | 0.036 | | | 2.614 | 0.003 | | | 1.000 | 0.000 | | |
| ***Plot:FD*** | 1.747 | 0.003 | | | 2.248 | 0.000 | | | 1.078 | 0.046 | | |
| ***Plot:TD*** | 2.389 | 0.041 | | | 2.419 | 0.000 | | | 1.000 | 0.022 | | |

| | *Parametric coefficients* | | | *Transect* | *Parametric coefficients* | | | *Transect* | *Parametric coefficients* | | | *Transect* |
|---|---|---|---|---|---|---|---|---|---|---|---|---|
| | *Estimate* | *Std. Error* | *p-value* | | *Estimate* | *Std. Error* | *p-value* | | *Estimate* | *Std. Error* | *p-value* | |
| ***Intercept*** | -0.442 | 0.110 | 0.000 | | 0.122 | 0.118 | 0.302 | | 0.538 | 0.129 | 0.000 | |
| ***FD*** | 0.177 | 0.155 | 0.256 | | 0.105 | 0.166 | 0.530 | | -0.829 | 0.182 | 0.000 | |
| ***TD*** | -0.479 | 0.155 | 0.002 | | 0.097 | 0.166 | 0.563 | | -0.942 | 0.182 | 0.000 | |
| | ***Smooth terms*** | | | 4 | ***Smooth terms*** | | | 5 | ***Smooth terms*** | | | 6 |
| | *edf* | *p-value* | | | *edf* | *p-value* | | | *edf* | *p-value* | | |
| ***Plot:SD*** | 1.001 | 0.000 | | | 2.294 | 0.000 | | | 1.000 | 0.020 | | |
| ***Plot:FD*** | 2.448 | 0.000 | | | 2.112 | 0.004 | | | 1.533 | 0.008 | | |
| ***Plot:TD*** | 2.366 | 0.012 | | | 1.839 | 0.000 | | | 1.000 | 0.272 | | |

**Table S3.** Generalized additive model (GAM) results for all β–diversity relations with the sea-inland gradient. Each box contains a single model output for the considered transect. Parametric coefficients are reported in the upper part of the box together with their estimate, standard error and p-value. β– SD was set as reference category for each model. A separate smooth function was performed for each facet of β–diversity. Smooth terms are reported in the lower part of the graph together with their effective degree of freedom (edf) and p- values.

| | *Parametric coefficients* | | | *Transect* |
|---|---|---|---|---|
| | *Estimate* | *Std. Error* | *p-value* | 1 |
| ***Intercept*** | 0.479 | 0.007 | 0.000 | |
| ***FD*** | 0.058 | 0.010 | 0.000 | |
| ***TD*** | 0.019 | 0.010 | 0.064 | |
| | ***Smooth terms*** | | | |
| | *edf* | *p-value* | | |
| ***PlotDist:SD*** | 3.600 | 0.000 | | |
| ***PlotDist:FD*** | 3.038 | 0.000 | | |
| ***PlotDist:TD*** | 3.165 | 0.000 | | |

| | *Parametric coefficients* | | | *Transect* |
|---|---|---|---|---|
| | *Estimate* | *Std. Error* | *p-value* | 2 |
| ***Intercept*** | 0.460 | 0.005 | 0.000 | |
| ***FD*** | 0.045 | 0.007 | 0.000 | |
| ***TD*** | 0.069 | 0.007 | 0.000 | |
| | ***Smooth terms*** | | | |
| | *edf* | *p-value* | | |
| ***PlotDist:SD*** | 2.344 | 0.000 | | |
| ***PlotDist:FD*** | 3.506 | 0.000 | | |
| ***PlotDist:TD*** | 1.108 | 0.000 | | |

| | *Parametric coefficients* | | | *Transect* |
|---|---|---|---|---|
| | *Estimate* | *Std. Error* | *p-value* | 3 |
| ***Intercept*** | 0.465 | 0.005 | 0.000 | |
| ***FD*** | 0.063 | 0.008 | 0.000 | |
| ***TD*** | 0.077 | 0.008 | 0.000 | |
| | ***Smooth terms*** | | | |
| | *edf* | *p-value* | | |
| ***PlotDist:SD*** | 2.648 | 0.000 | | |
| ***PlotDist:FD*** | 3.423 | 0.000 | | |
| ***PlotDist:TD*** | 1.001 | 0.000 | | |

| | *Parametric coefficients* | | | *Transect* |
|---|---|---|---|---|
| | *Estimate* | *Std. Error* | *p-value* | 4 |
| ***Intercept*** | 0.552 | 0.006 | 0.000 | |
| ***FD*** | -0.029 | 0.009 | 0.001 | |
| ***TD*** | -0.015 | 0.009 | 0.098 | |
| | ***Smooth terms*** | | | |
| | *edf* | *p-value* | | |
| ***PlotDist:SD*** | 1.001 | 0.000 | | |
| ***PlotDist:FD*** | 3.296 | 0.000 | | |
| ***PlotDist:TD*** | 1.000 | 0.000 | | |

| | *Parametric coefficients* | | | *Transect* |
|---|---|---|---|---|
| | *Estimate* | *Std. Error* | *p-value* | 5 |
| ***Intercept*** | 0.652 | 0.006 | 0.000 | |
| ***FD*** | -0.129 | 0.009 | 0.000 | |
| ***TD*** | -0.108 | 0.009 | 0.000 | |
| | ***Smooth terms*** | | | |
| | *edf* | *p-value* | | |
| ***PlotDist:SD*** | 3.701 | 0.000 | | |
| ***PlotDist:FD*** | 3.180 | 0.000 | | |
| ***PlotDist:TD*** | 3.260 | 0.000 | | |

| | *Parametric coefficients* | | | *Transect* |
|---|---|---|---|---|
| | *Estimate* | *Std. Error* | *p-value* | 6 |
| ***Intercept*** | 0.579 | 0.007 | 0.000 | |
| ***FD*** | -0.086 | 0.010 | 0.000 | |
| ***TD*** | -0.017 | 0.010 | 0.093 | |
| | ***Smooth terms*** | | | |
| | *edf* | *p-value* | | |
| ***PlotDist:SD*** | 1.000 | 0.000 | | |
| ***PlotDist:FD*** | 1.783 | 0.000 | | |
| ***PlotDist:TD*** | 1.000 | 0.000 | | |
